# Dynamic exometabolomics reveals metabolic adaptations of *Staphylococcus epidermidis* to pH-mimicking skin and bloodstream

**DOI:** 10.1101/2025.07.11.664416

**Authors:** Elisabete Morais, Ana M Gil, Maria Miragaia, Luís G Gonçalves, Ana V Coelho

## Abstract

*Staphylococcus epidermidis* (SE) is a common human skin coloniser, which is often the cause of medical device-associated infections. SE population is composed of two clonal lineages, A/C and B, with distinct pathogenic potential. Although pH is known to change during infection when SE crosses the host skin to access the bloodstream, the impact of this pH alteration on SE pathogenicity is poorly understood. Recognizing how SE deals with pH increments will help designing effective prevention and treatment strategies against SE infections. To investigate the metabolic adaptations of representative A/C and B strains to different pH, we mimicked skin and blood pH conditions (5.5 and 7.4) and followed biomass formation, growth media pH and exometabolites over time. Although both strains share some metabolic patterns, specificities were identified for each strain and pH condition. The B strain was better adapted to use diverse carbon sources and at blood pH has a more active TCA cycle and amino acid catabolism. At blood pH, the B strain depletes formate from the extracellular media, while its extracellular accumulation by the A/C strain could work as a host invasion strategy. For both SE strains, TCA cycle regulation, purine biosynthesis and glutamate uptake could be associated with virulence, particularly biofilm production, especially relevant for ICE25 which is able to produce high adherence biofilm. The uptake and consumption of saccharides follow similar profiles and seem to be pH-regulated by both strains. The dynamic study of SE exometabolome has contributed to understanding the intracellular processes and their relationship with virulence.

## Introduction

*Staphylococcus epidermidis* (SE), the main human skin coloniser, is the major cause of nosocomial infections associated with the use of medical devices^1,2^. A recent study reported that 80% of adults admitted to Mayo Clinic Rochester (USA) between January 2016 and July 2022 with monomicrobial coagulase-negative staphylococci bloodstream infections are SE positive^3^. Moreover, during a Belgian national surveillance, it was determined that 77% of SE infections were resistant to methicillin^4^. These facts justify the urgent need for a better understanding of this bacteria’s virulence mechanisms so that emerging infections may be prevented and treated more efficiently.

SE is an opportunistic pathogen that presents a dual behaviour of commensalism and pathogenicity^5,6^. SE isolates were phylogenetically classified as belonging to A/C or B lineages^7,8^. While the first lineage equally comprises strains with pathogenic and commensal potential, the second lineage is mostly populated by strains collected from colonised individuals^7,8^. Despite colonising mainly the human skin and nares, SE can be present in different human body microenvironments^6^. At such sites, bacteria are exposed to different oxygen, temperature, salinity, lipid content, osmolarity and pH, among other conditions^9^. Several distinct conditions are also experienced when SE accesses the bloodstream, for instance *via* medical devices, often triggering an infectious process. The most important SE virulence factor is its ability to form biofilms attached to abiotic surfaces^10,11^. However, the process explaining why SE strains belonging to A/C clonal lineage have a higher pathogenic potential is not fully understood. Méric *et al*. suggested a mechanism involving the horizontal gene transfer of pathogenic determinants allowing an adaptation into the invasive niche^12^. No study has ever compared the metabolism of SE strains belonging to clonal lineages A/C and B. Only a few targeted studies reported their differences at the gene and protein levels. Yang *et al*. found that two proteins were highly correlated with a pathogenicity profile, a targeting RNAIII-activating protein and the accumulation associated protein (AAP)^13^. Interestingly, it was also suggested that the significant difference between the ratio of non-identical *vs*. identical genes coding for surface proteins from the translation-related genes and total genes could be related with pathogenicity^14^. Moreover, differences among SE strain-specific secreted antigens and adhesins could explain the higher virulence capability of some pathogenic strains^15^.

Moreover, only a scarce number of metabolic studies involving SE strains were published and they rely on a snapshot of metabolic information not fully demonstrating the dynamics of SE metabolism during bacterial growth. However, several environmental stressors have been shown to inactivate the tricarboxylic acid (TCA) cycle with intermediate metabolites redirected into polysaccharide intercellular adhesin (PIA) synthesis, a key constituent of biofilms^16,17^. Recently, based on the study of the intracellular proteome and metabolome, we described the metabolic adaptations of strains belonging to A/C and B lineages to pH-mimicking skin and blood conditions at the mid-exponential phase^18^ (unpublished data). Therefore, we hypothesised that the adaptation of each representative strain to each pH also induces differences on their time course exometabolome profiles reflecting the diverse intracellular metabolic processes reported^18^. Consequently, characterization of the exometabolome will contribute with complementary information to understand the differential impact of media pH on the pathogenic capacity of *S. epidermidis* strains. In particular, may help to identify metabolites or mechanisms that explain why a strain takes advantage to cause host infection at pH 7.4 and/or to proliferate at skin pH. We selected representative strains of each one of the clonal lineages, A/C (ICE25 strain) and B (19N strain), and grew them at skin and blood pH-mimicking conditions. Optical density, media pH and exometabolome Nuclear Magnetic Resonance (NMR) profile were monitored during cellular growth. Understanding how SE A/C or B strains adapt to abiotic factors, such as media pH, is of major importance in revealing the mechanisms underlying the dual behaviour of this bacterial species^6^.

## Results and Discussion

To investigate the dynamic adaptation of 19N and ICE25 strains to skin (pH 5.5) and blood (pH 7.4) pH conditions, we analysed their extracellular metabolomes using ^1^H-NMR. Unexpectedly, during the analysis of these data, we observed significant differences in the concentrations of 20 metabolic features prior to inoculation (t_0h_), between the media at pH 5.5 and pH 7.4, with variations ranging from −80% to 70% (Supplementary Fig. S1). For example, the concentration of glucose, a predominant metabolite in the medium and a primary carbon source known to suffer pyrolysis during autoclaving^19^, was 29% higher in the medium at pH 5.5 than at pH 7.4. This difference in the initial glucose levels was attributed to the sterilisation procedure, glucose levels decreased by 6% and 25% relative to pre-autoclaved TSB, respectively, for pH 5.5 and pH 7.4 (Supplementary Fig. S2). To assess whether these initial differences influenced bacterial metabolism during growth we compared the metabolome of five ICE25 biological replicates harvested at mid-exponential phase in both autoclaved and filtered-sterilized media (Method description in Supplementary Information). Multivariate analysis revealed significant differences in metabolites concentrations only between the two pH conditions, irrespective of the sterilization method (Supplementary Fig. S3).

These findings highlight the importance of selecting a sterilisation method, such as filtration, that preserves medium composition independently of pH, as medium alterations may impact bacterial growth^20^. Nevertheless, in this study, we focus on inter-strain comparisons conducted under the same pH conditions. Comparisons between different pH environments take into account the initial differences in metabolite concentrations observed at t_0h_.

### Growth rate is similar between commensal and pathogenic strains

The measurements of OD_600nm_ and medium pH for the original and adjusted growth periods are presented in Supplementary Figure S4 and Figure 1, respectively. Growth parameters were calculated using the OD_600nm_ measurements in Supplementary Table S1. No significant differences in growth rates are found between strains at the same pH conditions (Fig. 1, Table 1). However, at pH 7.4, 19N reaches significantly higher biomass than ICE25 (OD_600nm_ of 10.93 ± 0.15 and 9.60 ± 0.14 for 19N and ICE25, respectively). Although these data suggest that both strains are metabolically adapted to a pH environment mimicking blood infection, this effect is slightly more noticeable for 19N. At pH 5.5, OD_600nm_ does not rise above 7 for both strains.

**Table 1.** SE growth parameters and extracellular pH for each experimental condition. Values correspond to average ± standard deviation. Student’s t-test was performed between strains within each pH condition (*** p-value < 0.001; ** p-value < 0.01). pH_0h_, pH_12h_, pH_min_ correspond to pH values at t=0h and t=12h and to the minimum pH value for each growth condition, respectively

|  | pH 5.5 |  | pH 7.4 |  |
| --- | --- | --- | --- | --- |
|  | 19N | ICE25 | 19N | ICE25 |
| Growth rate at exponential phase ($h^{-1}$) | $0.87 \pm 0.14$ | $0.80 \pm 0.04$ | $0.90 \pm 0.02$ | $0.89 \pm 0.07$ |
| Maximum OD | $6.75 \pm 0.12$ | $6.57 \pm 0.85$ | $10.93 \pm 0.19$ | $9.61 \pm 0.14^{***}$ |
| Minimum pH ($pH_{min}$) | $4.71 \pm 0.05$ | $4.77 \pm 0.10$ | $5.82 \pm 0.05$ | $6.12 \pm 0.02^{***}$ |
| pH decrease $pH_{0h} - pH_{min}$ | $0.91 \pm 0.09$ | $0.88 \pm 0.07$ | $1.33 \pm 0.03$ | $1.05 \pm 0.03^{***}$ |
| **pH increment $\text{pH}_{12\text{h}} - \text{pH}_{\text{min}}$** | $0.21 \pm 0.09$ | $0.26 \pm 0.16$ | $1.09 \pm 0.14$ | $0.65 \pm 0.06^{**}$ |

**Figure 1.**
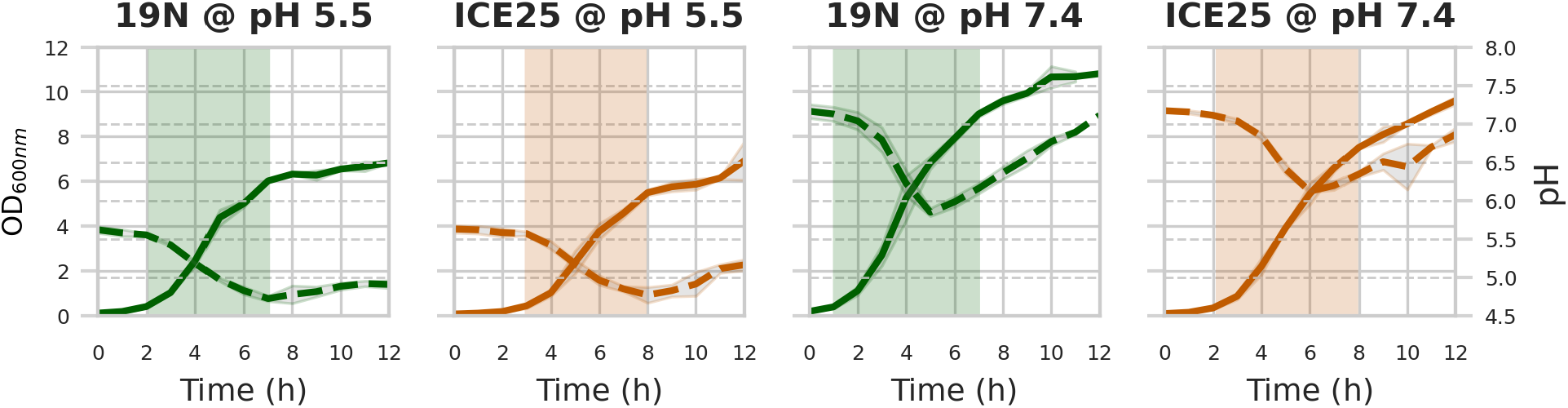
Extracellular pH recovers only at blood-mimicking conditions after SE acidification of the media during exponential growth. Biomass (OD_600nm_, optical density at 600 nm, left y-axis, solid grid line) and extracellular pH (right y-axis, dashed grid lines) average curves for 19N and ICE25 strains grown in TSB at initial pH of 5.5 and 7.4 (n=3). Time was adjusted as described in the methods section. Filled and dotted lines correspond to OD_600nm_ and extracellular pH measurements, respectively, with standard deviation in light grey. The shaded areas correspond to the exponential phase for each condition.

### Exometabolic profiles differ between strains at the same initial pH

A total of 78 metabolic features, 47 assigned metabolites (comparable to the number of metabolites identified by NMR in other extracellular media from bacteria/staphylococcus^28^) and 31 unknowns (Unks) were quantified following NMR spectra analysis (Supplementary Table S2 and Supplementary Fig. S4). Identification of these Unks should be accomplished in further studies, at least for those showing differential patterns between strains at the same pH (eg. Unk_5.45, Unk_5.94 and Unk_x). All metabolites were detected in the four experimental conditions, except for ethanol not detected at pH 5.5 for ICE25 and *cis*-aconitate at the same pH for both strains. To evaluate temporal variation for each metabolic feature in each experimental condition, a permutation filter analysis was performed, as previously described^21^. This approach showed that at pH 5.5, the levels of 64 and 59 metabolic features change respectively during 19N and ICE25 growth, and at pH 7.4, 49 and 68 metabolic features suffer alteration during the growth of 19N and ICE25 (Supplementary Table S4). Interestingly, ICE25 at pH 7.4 exhibits the highest number of varying features. For both strains at pH 5.5, only tryptophan and trehalose concentrations do not change; at pH 7.4, the same happens for lysine and tyrosine. For each strain, some metabolites maintain their levels at both pH conditions, namely citrate, raffinose and trehalose for 19N and isoleucine for ICE25. Only the metabolic features with temporal variations were considered for further analyses.

Moreover, cluster analysis addressed whether the same metabolic feature presents a similar temporal profile for both strains within the same pH. Three and six clusters were determined respectively, for pH 5.5 and 7.4 (Fig. 2a and 2d, Supplementary Table S5). For pH 5.5, an uptake (decrease along time), a secretion (increase along time) and a mixed (combination of increasing and decreasing periods) cluster were identified (Fig. 2a). While the uptake cluster is more populated by metabolic features measured for ICE25, the mixed cluster is mainly populated by 19N metabolic features. For pH 7.4, metabolic features are clustered in fast and slow uptake, fast and slow secretion, and two mixed profiles (Fig. 2d). Slow-uptake, slow and fast-secretion clusters integrate mainly ICE25 metabolic features, while the fast-uptake cluster includes more 19N metabolic features. Some features are common to both uptake profiles but consumed at different rates by each strain, this is the case of leucine, mannose, serine and the Unk_3.69 that were found to be uptake faster by 19N than by ICE25. Although both strains at pH 5.5 secreted ornithine, this cluster analysis points out that the secretion pattern over time significantly differs between strains (secretion and mixed profiles, respectively, for 19N and ICE25). At pH 7.4, formate is part of the slow-uptake cluster for 19N, while it is included in the slow-secretion cluster for ICE25.

**Figure 2.**
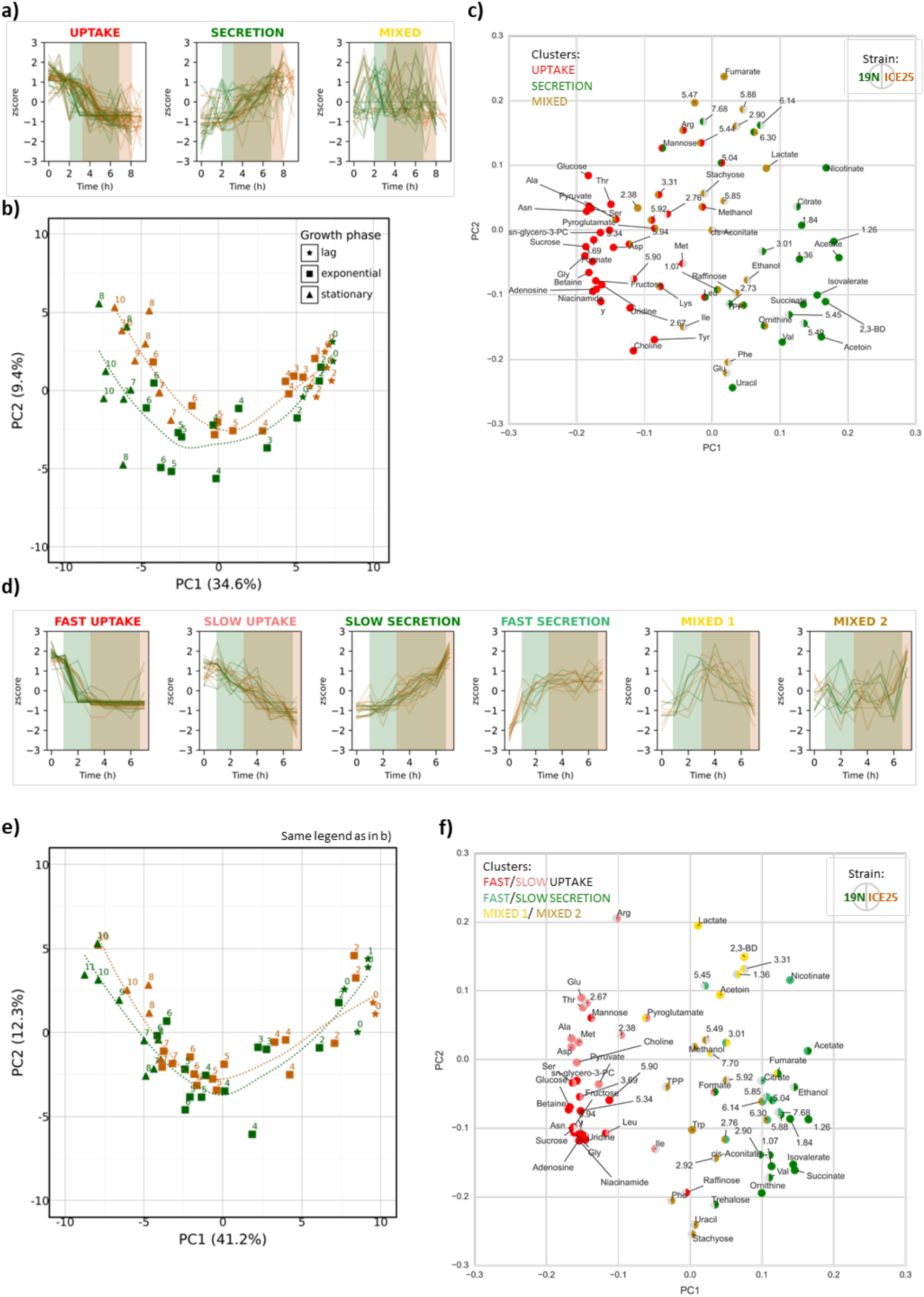
Time course metabolite clustering and PCA analysis for 19N and ICE25 grown at pH 7.4 and 5.5. K-means clustering with dynamic time warping of the metabolic features profiles measured for each strain at pH 5.5 (A) and pH 7.4 (D). Shaded areas correspond to the exponential phase defined in figure 1. PCA scores plots of extracellular metabolites over 19N and ICE25 growth at pH 5.5 (B) and 7.4 (E). For K-means clustering and PCA analysis, metabolic feature concentrations were mean centred and divided by standard deviation (z-scores). 19N and ICE25 exponential phase are represented by green and orange shaded areas, respectively. Dotted lines represent the trajectory of growth for each strain within each pH. PCA loadings plots (C, F) represent the metabolic features weight of each component in the respective PCA scores plots (B, E). Each metabolic feature is coloured by the cluster they belong in 19N (left half circle) and ICE25 strain (right half circle). For visualization purposes, some metabolic feature names are abbreviated (see Supplementary Table S2).

PCA analysis was performed for each pH condition considering both strains and all the metabolic features during the entire time course. The obtained patterns show a sequential grouping of samples over time evidencing the several growth phases along principal component 1 (PC 1) and a differentiation of the exponential relative to the other two phases (PC2) (Fig. 2b and 2e). PCA loadings plots (PC1 vs PC2) for both strains (Fig. 2c and 2f) highlight, as expected based on the PCA scores-plots, that uptake metabolites are increased at initial incubation times, while the secreted ones are increased at later growth times. Both types of metabolites are responsible for the separation in PC1. Furthermore, the mixed profiles are responsible for the separation in PC2, which mainly change in the exponential phase.

### Saccharides uptake is regulated by media pH

According to our data, both SE strains preferentially utilise monosaccharides such as glucose, mannose, and fructose, as well as the disaccharide sucrose (Fig. 3a), even in the presence of other oligosaccharides like trehalose, stachyose, and raffinose (Supplementary Fig. S5). The extracellular levels of trehalose and raffinose do not change during 19N growth and those of stachyose for both strains at pH 5.5 (Supplementary Table S3). SE ATCC12228 strain has grown when glucose was replaced by raffinose or trehalose in synthetic minimal medium^22^, suggesting that at least this strain has the mechanisms to uptake these oligosaccharides. In this same study, SE ATCC12228 could not use mannose as a single carbon source, which was not the case for 19N and ICE25. Previous research has shown that SE can oxidise this monosaccharide^23^. At an initial pH of 5.5, media was first depleted in fructose, followed by sucrose, glucose, and mannose (only completely depleted for 19N, Supplementary Fig. S4). However, at pH 7.4, the three first saccharides were depleted simultaneously followed by mannose, indicating an earlier and higher uptake of carbon sources (Fig. 3a). Depletion of glucose or mannose from the media matches with the end of the exponential phase, respectively at pH 5.5 and 7.4. 19N when grown at blood pH, was previously shown to present several elevated PTS protein levels when compared with skin pH, including fructose-specific IIABC component (SERP2260), sucrose-specific IIBC component (SERP1968), and glucose-specific EIICBA component (SERP2114)^18^. This supports the fact that at pH 7.4, there is a higher demand for carbon sources, which may be associated with the higher biomass that was measured at the stationary phase, especially for 19N.

**Figure 3.**
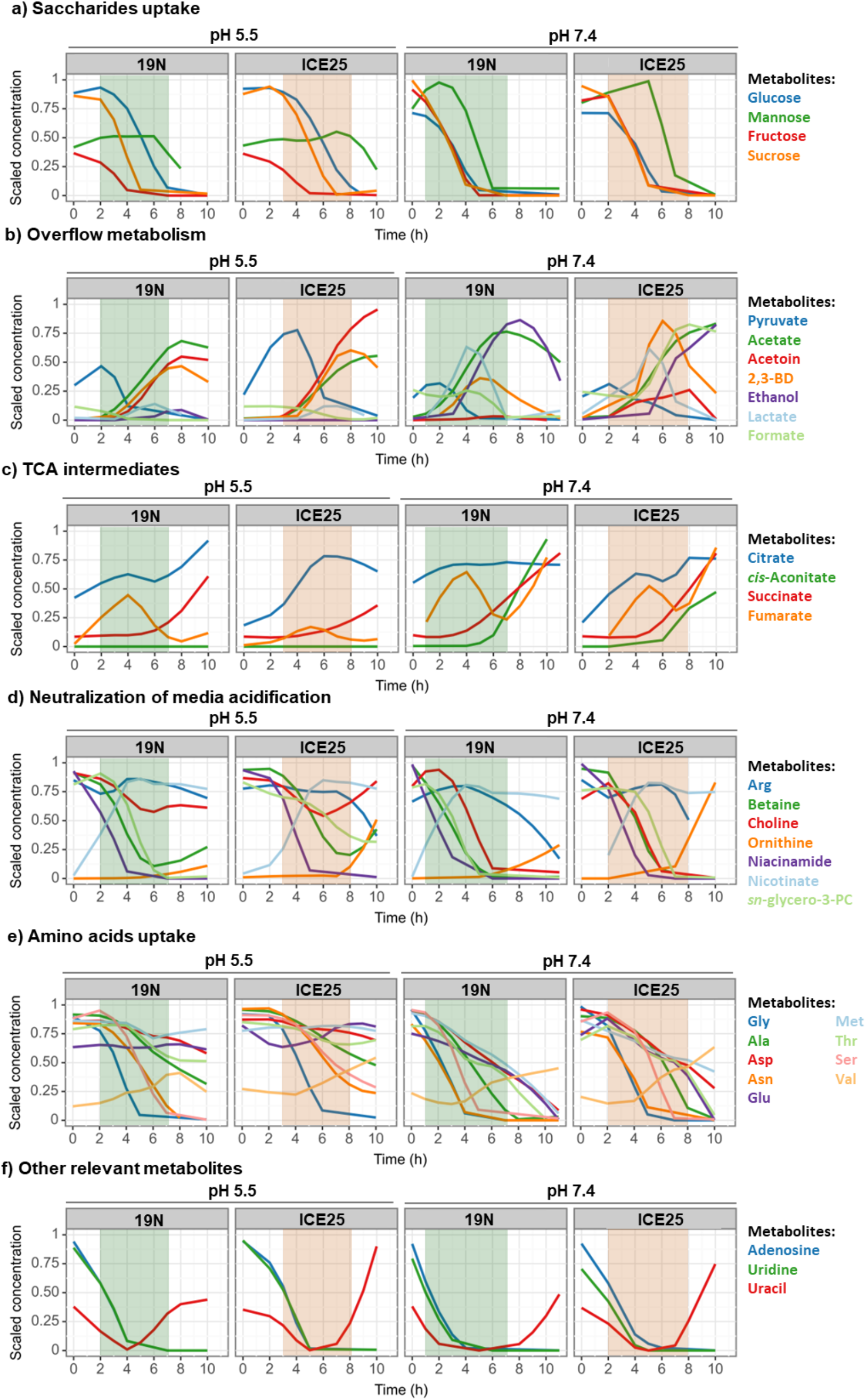
Exometabolome adaptations of 19N and ICE25 strains to both skin and blood mimicking pH conditions. In each panel are the scaled metabolite profiles depicted in the respective Results and Discussion chapter. For each metabolite, concentrations are scaled to range from zero to one (preprocessing.MinMAxScaler function used from Scikit-learn 1.4.1). Note that comparisons between concentrations of different metabolites are not possible. Only comparisons between conditions for each metabolite profile can be done. Shaded areas correspond to the defined exponential phase in figure 1.

It is noteworthy that, despite their different starting concentrations (fructose concentration was 93% less than that of glucose), fructose uptake was favoured over glucose at acidic pH levels (Fig. 3a). The genomes of both SE strains encode a PTS fructose-specific EIIABC component (SERP0359), which sustains fructose uptake by phosphorylating it to fructose-1-phosphate. Subsequently, this compound can be further phosphorylated by 1-phosphofructokinase (PfkB) to fructose-1,6-bisphosphate, which is redirected to glycolysis. This path can be considered a shortcut because it relies on one reaction less than when glucose is uptake for glycolysis. Mild acid stress caused to *S. aureus* showed that the transcription of a protein similar to the PTS fructose-specific enzyme IIBC component was upregulated in the first 3 h post-stress^24^, suggesting that *Staphylococcus* species grown at acidic pH tend to use fructose as the first-choice carbon source.

Apart from the transporter mentioned above, both strains can also internalise glucose using a glucose uptake protein (GlcU, SERP1838) and mannose through a putative PTS system mannose-specific EIIBCA component (SERP2260). The latter has been suggested as a multi-stress responder in SE^25^.

None of the strains seems to be very selective for glucose uptake within the pool of saccharides, contrary to what was previously suggested for *S. aureus* at pH 7.4^26^. Overall, the consumption of monosaccharides and sucrose is probably regulated at some level by extracellular pH for both strains. Most of these saccharides feed glycolysis and, subsequently, can feed the overflow metabolism or TCA cycle, depending on the nutritional and environmental conditions.

### Overflow metabolism occurs to a higher extent at blood pH for both strains

Overflow metabolism is known in several bacterial species and occurs under glucose excess conditions leading to reduced carbon flux into the TCA cycle due to carbon catabolite repression (CCR), and consequently limiting oxidative phosphorylation, and up-regulating glycolysis with the secretion of several by-products^30^. NADH molecules from glycolysis and the pyruvate dehydrogenase complex provide electrons to the electron transport chain, which drives ATP synthesis through oxidative phosphorylation and regenerates NAD+, crucial for the continuity of glycolysis and pyruvate dehydrogenase complex reactions. Although overflow metabolism produces less ATP per glucose molecule than respiration does (2 ATP vs 22 ATP), it is proposed to be preferred at high growth rates due to its lower protein biosynthesis cost^31^. Moreover, when glucose is depleted, a shift to TCA occurs and acetate, originated from acetyl-coenzyme A by the Pta/Acka pathway (acetate kinase (Ack)-phosphate acetyltransferase (Pta)) during overflow metabolism, can be uptake and oxidized via the TCA cycle.^27^ Our data support that during exponential growth, acetate is accumulated in the extracellular media in high concentration showing that SE is resorting to the Pta/Acka pathway. After, in the stationary phase, occurs a shift to TCA and the levels of acetate stabilize and decrease. In the present study, besides acetate several other fermentation products originated from pyruvate were quantified: lactate, acetoin, 2,3-butanediol (2,3-BD) and ethanol (Supplementary Table S2). At both pH conditions, pyruvate was accumulated in the media during the lag phase, being this situation extended to the early exponential phase for ICE25 at pH 5.5 (Fig. 3b). Its final uptake matches the starting of acetoin accumulation at acidic pH, lactate and ethanol at pH 7.4, and acetate and 2,3-BD at both pHs. A similar variation in pyruvate levels was previously reported for *S. aureus*^28,29^. The use of pyruvate in more catabolic processes at pH 7.4 can explain its higher accumulation at acidic pH.

Our previous studies on the intracellular differential metabolome and proteome at the mid-exponential phase have shown that at pH 7.4, both SE strains resort to overflow metabolism to a greater extent than at pH 5.5^18^. These results led to the hypothesis that cellular energy is required at a higher rate at blood pH. According to these results, acetate is the main product of the overflow metabolism at both pH, followed by lactate at pH 7.4 (Supplementary Fig. S5). Acetate was also found to be the most prominent overflow metabolite of *S. aureus* grown in RPMI (pH 8.2), among lactate, 2-acetolactate, acetoin and 2,3-BD^28^. Acetoin is the only fermentation product with lower levels at pH 7.4 for both strains (Supplementary Fig. S5). Higher levels of acetoin and 2,3-DB were measured for ICE25 at both pH, while 19N at pH 7.4 does not secrete acetoin (Fig. 3b), suggesting that acetoin synthesis is highly plastic to pH and strain dependent. Accumulation of acetoin and 2,3-BD was reported before for SE biofilms under anaerobiosis^15^. Common temporal profile variations at pH 5.5 were found for the extracellular levels of acetate, acetoin and 2,3-DB for both strains, whose levels increase were followed by none or reduced uptake at the stationary phase (Fig. 3b). At acidic pH, both strains secrete residual amounts of lactate and ethanol, being lactate fully consumed before the end of the assay (Fig. 3b).

At pH 7.4, lactate, acetoin and 2,3-DB uptake at later growth phase occur for both strains, while 19N demonstrates extra potential to uptake acetate and ethanol from media at the stationary phase (Fig. 3b). Ethanol extracellular levels start to increase at the beginning of the exponential phase for 19N and slightly later for ICE25 (Fig. 3b). Lactate follows a very similar temporal pattern for both strains, with its levels higher around the mid-exponential (Fig. 3b). As reported for *S. aureus* at pH 7.0^32^, when glucose is at low levels, its growth is sustained by aerobic utilisation of lactate rather than using the overflow acetate. Probably at pH 7.4, until the mid-exponential, pyruvate is reduced to lactate by lactate dehydrogenase (Ldh, SERP2156), thus leading to increased lactate intracellular levels and, consequently, its extracellular levels. The latter starts to decrease when extracellular saccharides are consumed and are converted by Ldh to pyruvate, generating reducing power used for oxidative phosphorylation by a lactate:quinone oxidoreductase. This lactate translocation between intra and extracellular compartments can be mediated by an L-lactate permease (SERP1957), which is known to be present in both strain’s genomes. Despite a non-preferred carbon source, acetate starts to be internalised by 19N when lactate is almost completely used up and is anticipated relative to ICE25 at blood pH (Fig. 3b). *S. aureus* imports and oxidises acetate only in the post-exponential phase, forming acetyl-CoA to enter the TCA cycle^26,29^. Moreover, the expression of sigma factor B regulator protein (rsbU, SERP1680) was suggested as a requirement for acetate catabolism^33,34^. This gene is present in the genome of both SE strains under study.

A more active overflow metabolism at pH 7.4 could be related to the higher initial levels of some saccharides and their faster depletion (Fig. 3a). Higher secretion of neutral overflow metabolites (acetoin and 2,3-BD) at acidic pH is very likely induced by media initial pH difference. Overall, the overflow metabolism differences between the two SE strains at each pH condition rely mainly on temporal adjustments for the secretion/uptake of certain metabolites rather than on major profile differences.

### TCA cycle is more active at pH 7.4 for 19N

In a nutrient-rich environment, only a low amount of pyruvate is diverted to the TCA^34^. Instead, it is catabolized by PTA/AckA pathway, but when preferential carbon sources decrease below the critical level, pyruvate catabolism is enhanced through TCA cycle to maintain bacterial growth^34^. A time-resolved study following the glucose excess-to-starvation transition in *S. aureus* at pH 7.0 has shown the accumulation of TCA cycle enzymes together with an increase of its extracellular levels^29^. Altogether, a similar variation pattern for the extracellular TCA cycle metabolites was observed for the two SE strains. At pH 5.5, we have determined that when saccharide starvation approaches (Fig. 3a), an extracellular accumulation of succinate starts for both strains and of citrate and fumarate for 19N (Fig. 3c). At blood pH, there is a higher extracellular accumulation of TCA cycle intermediates relative to skin pH. In addition to fumarate and succinate, *cis*-aconitate also accumulates at the late-exponential phase, mainly for 19N (Fig. 3c). A decrease of overflow metabolites (acetate, lactate, 2,3-BD or ethanol) (Fig. 3b) also shows that TCA cycle is activated at the late-exponential, promoting their catabolism. Lactate and 2,3-DB are consumed by both strains at the two pHs generating NADH, while acetate and ethanol were only consumed by 19N until the end of our assay. The conversion of acetate to acetyl-CoA requires one ATP, which could be the reason for a preferred consumption of lactate and 2,3-BD.

During the early exponential, citrate levels only increase for ICE25 (Fig. 3c). In contrast, both strains accumulate fumarate at the two pHs, however, in a modest amount by ICE25 at pH 5.5. Fumarate is consumed when saccharide levels decrease (Fig. 3c) and may be used as a carbon source after saccharide depletion.

SE TCA cycle is important to respond to multiple stressors^16,35^, and to regulate virulence factors synthesis and antibiotic resistance^34^. TCA cycle derepression in *S. aureus* coincides with the maximal expression of some secreted virulence factors^26^. Future determination of these factors’ levels and their correlation with both SE strains growth phases should be done. TCA cycle plasticity, which is corroborated by our data showing its pH, strain and growth phase-dependency, allows the creation of diverse strategies of pathogenicity associated with different metabolic situations. (This topic is further discussed in chapter “Regulation of Carbon Central Metabolism and Virulence factors”).

### Neutral and vitamin B3-related metabolites, quaternary ammonium compounds, and ammonia release counteract acetate overflow at blood pH

For all experimental conditions, media acidification was found during bacterial growth (Fig. 1 and Supplementary Table S1). At pH 5.5, a minimum pH of around 4.7 was reached for both strains at the late-exponential (7h and 8h, respectively, for 19N and ICE25), after which it tended to increase slightly (Fig. 1, Table 1). When simulating blood pH, the minimum pH reached at 6 h for both strains is significantly lower for 19N (5.82 ± 0.10 and 6.12 ± 0.02 for 19N and ICE25, respectively). At this condition, 19N showed a good ability to recover initial pH values, reaching 0.1 pH units above ICE25. For *S. aureus* also grown in TSB, medium acidification was reported at pH 6 matching with the post-exponential phase, followed by a pH increase to 8, at the stationary phase^26^. In our experiments this phase was not fully reached, a pH trend increase over time could happen for both strains (Fig. 1).

The extracellular pH decrease observed is most likely due to secretion of acidic compounds and subsequent medium pH increase, at condition pH 7.4, is partially due to their uptake. Strong linear correlations (>|0,8| and *p*-value <0.05) were found between pH and the levels of some metabolites that can directly impact extracellular pH at the exponential phase in all conditions, and mainly during the post-exponential for 19N at pH 7.4 (Fig. 4, Supplementary Fig. S6). For pH 5.5, acetate and nicotinate secreted levels correlate negatively with pH decrease for both strains and those of isovalerate and succinate, respectively, for 19N and ICE25. Betaine and choline, two positively charged quaternary ammonium compounds, correlate positively with this pH decrease for both strains and only for 19N, respectively. At pH 7.4, secretion of acetate by both strains, lactate, nicotinate and fumarate by ICE25 and consumption of choline and betaine by both strains are associated with media acidification. The uptake of acetate and the secretion of ornithine by 19N seem to participate in the recovery of media pH.

**Figure 4.**
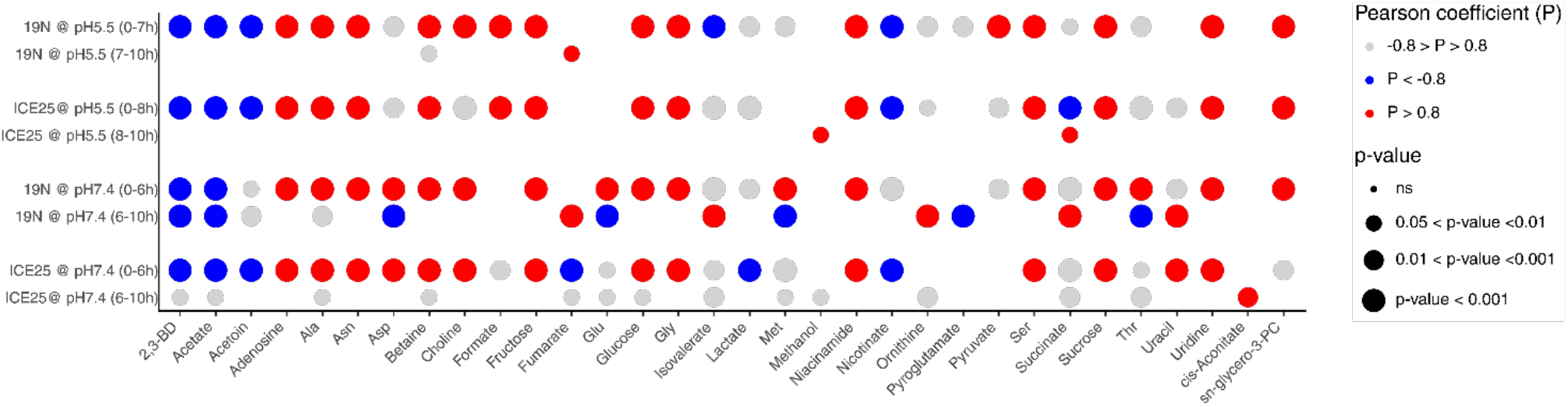
Correlation of metabolite profiles with pH dynamics depends on the experimental condition. Metabolic feature correlation with extracellular pH for each experimental condition and each pH variation. Pearson correlation coefficient was determined as their associated *p*-values. This figure represents the metabolic features with a strong and statistically significant correlation with pH (>|0,8| and p-value <0.05). Some metabolic feature names are abbreviated (see Supplementary Table S2). The results of all the metabolic features are detailed in the Supplementary Figure S5.

Although several metabolites may contribute to media acidification during the exponential phase, we attribute this effect mainly to acetate due to its higher accumulation (10 mM and 15 mM, respectively, at pH 5.5 and 7.4). At pH 7.4, lactate is the second major contributor reaching an extracellular concentration around 4 mM (Supplementary Table S2). After an initial acetate overflow, neutral compounds (acetoin and 2,3-BD) start to be secreted (Fig. 3b), thus helping to prevent excessive media acidification, matching what was previously reported^36^. The secretion of these metabolites is reduced when media acidification is reverted, except for acetoin for ICE25 at pH 5.5 (Fig. 3b). At this point, acetate secretion rate is also reduced or its uptake starts, respectively for ICE25 and 19N (Fig. 3b). Besides, at pH 7.4, any of these compounds is exported (Fig. 3d). Instead, ornithine (most likely co-transported by arginine/ornithine antiporter, ArcD, SERP2247) is accumulated in the media mainly by ICE25 (Fig. 3d). Ornithine is an intermediate of the urea cycle, which produces ammonium that can counteract extracellular acidification. Its accumulation has been previously reported in *S. aureus*^29^.

At each pH, both strains showed similar temporal extracellular profiles of choline and its related metabolites, betaine and *sn*-glycero-3-phosphocholine (Fig. 3d). At blood-mimicking pH, these metabolites are completely depleted from the media during the exponential phase, while at skin pH, only *sn*-glycero-3-phosphocholine is completely depleted from the media by 19N at the post-exponential phase (Fig. 3d). Then they are accumulated, suggesting that both metabolites start to be exported to help with media acidification control. Betaine and its precursor, choline, are associated with osmoprotective and pH homeostasis roles in SE^37^ and *sn*-glycero-3-phosphocholine is associated with glycerolipids metabolism and poly(ribitol phosphate) wall teichoic acid biosynthesis^37^. Although at pH 7.4 there is a more extensive import of betaine and choline, their intracellular levels at the middle-exponential are not enhanced relative to pH 5.5 for both strains^18^. These results suggest the existence in SE of betaine catabolism and choline synthesis pathways which have not yet been described. Betaine medium deprivation, concomitant with equivalent or low intracellular levels of choline and betaine occurring at pH 7.4 for both strains ^18^, is a puzzle that needs to be solved.

Under both pH conditions and strains, there is a strong correlation between the profiles of vitamin B3-related compounds, niacinamide and nicotinate, which we associate with the direct conversion of niacinamide into nicotinate (Fig. 3d). Niacinamide levels decrease from time zero reaching the depletion at 4h-6h for both strains and pH conditions. In contrast, nicotinate levels increase, reaching their maximum around 4h followed by a slow consumption. Niacinamide conversion to nicotinate is catalysed by a nicotinamidase (SERP1454), a homologous existing in both strains, leading to ammonium production. This reaction is part of the NAD salvage pathway, responsible for NAD^+^ and NADP^+^ synthesis. However, present results show that niacinamide import occurs at the same rate as nicotinate export, suggesting that all niacinamide imported is converted to nicotinate intracellularly and immediately secreted, not feeding the NAD salvage pathway and, probably, mainly contributing to reducing media acidification, which could be critical for SE survival. The exact reason for exporting nicotinate back to the media remains unclear.

Altogether, our experiments show that both strains can prevent a strong media acidification, mainly attributed to acetate secretion, and also, at pH 7.4 to lactate, thus avoiding reaching pH below 4.5 at which their growth rate is significantly reduced^38^. At pH 5.5, secretion of neutral compounds, (acetoin and 2,3-BD) and of betaine and choline contribute to this acidification control that could also be attributed to ammonium release during amino acid catabolism^26^ and to niacinamide/nicotinate conversion. Moreover, ICE25 resorts to the urea cycle to hamper extracellular acidification. At pH 7.4, extracellular pH is almost recovered by both strains being the ammonium producing pathways relevant in acidification control. Although 19N seems less efficient at controlling media pH decrease (Table 1), it manages a more effective recovery of initial blood pH.

### The arginine deiminase pathway is more active for the A/C strain

The arginine deiminase (ADI) pathway is involved in the acidic stress response of multiple bacteria, including SE^39^. Arginine uptake by the arginine/ornithine antiporter (ArcD, SERP2247) occurs with simultaneous ornithine export. Then, intracellular arginine can be converted to ornithine and ammonia by the arginine deiminase (ArcA, SERP2250). It has been demonstrated that the synthesis of NH_3_ via ADI, and its subsequent protonation generating NH_4_^+^, increases both intracellular and extracellular pH^39^. Indeed, in this study, a decrease in extracellular arginine levels was measured for both strains at both pH conditions starting at the post-exponential phase. Concomitantly, ornithine extracellular levels increased. This effect is more pronounced for ICE25 (Fig. 3d). The presence of the arginine catabolic mobile element (ACME), which comprises genes involved in the ADI pathway (among others), has been associated with SE strains with a higher pathogenicity potential^38,40^. An extra copy of arcD was found in ICE25 ACME and identified in its proteome (unpublished results), which explains the observed higher export of ornithine. Moreover, reduction of arginine levels can also be promoted by arginase activity to subvert the antimicrobial activity of macrophages that use arginine to obtain nitric oxide^41^.

### The A/C strain accumulates extracellular formate at blood pH

Formate levels at blood pH present a strain-specific profile. For the 19N, the residual formate is depleted from the media, while for ICE25, it accumulates extracellularly onward from the early-exponential phase (Fig. 3b). Only strains belonging to B lineage, populated by clinical isolates from healthy donors like 19N, possess a formate dehydrogenase (Fdh)^38^. This membrane-bound dehydrogenase has been associated with formate detoxification and NAD^+^ reduction in *S. aureus*^42^. For the 19N strain, independently of the initial pH, the residual formate present in the media seems to be oxidised by Fdh to CO_2_ at the membrane level by providing electrons to feed the menaquinone pool. Despite the A/C strain not relying on any known Fdh activity, formate was slightly consumed at skin pH. A formate channel (FocA) from *E. coli* (P0AC23) imports formate at pH<6.8^43^. A 70% similarity of ICE25 and 19N nitrite transporter (NirC) with FocA from *S. aureus*, annotated as formate/nitrite transporter, was calculated. Therefore, formate formed during the overflow metabolism does not seem to be exported since it does not accumulate at pH 5.5, but is exported at blood pH during the post-exponential phase (Fig. 3b).

The determined formate accumulation at blood pH may play a role in SE pathogenicity. Previous data showed that formate can have an impact on pathogenicity, either by protecting bacteria from antimicrobials ^44^ and from the host immune systems^45^, or by enhancing a pathogenic phenotype^45,46^.

### Amino acid catabolism is enhanced at blood pH

A total of seventeen amino acids were quantified in this study. All these compounds changed their levels between at least two experimental conditions (Supplementary Table S3). The predominant variation pattern is their consumption over time (Fig. 3d and Supplementary Fig. S5). At pH 5.5, the levels of six amino acids do not change for ICE25, one of them, tryptophan, also stays constant for 19N, being the only amino acid not changing for this strain. Two amino acids, lysine and tyrosine, do not change for both strains at pH 7.4 and other two only for 19N, phenylalanine and valine. Isoleucine maintains its levels for ICE25 at both pHs (Supplementary Table S3).

Valine is the only amino acid for which the levels increase over time (Supplementary Table S2). This behaviour occurs for all experimental conditions except for 19N at pH 7.4. The same pattern of secretion of valine is reported for *S. aureus* in TSB at pH 7.4^47^ and in RPMI^28^, being suggested that this amino acid could function as a quorum sensing involved in the development and maintenance of biofilm^47^.

*S. aureus* overcomes saccharide deficiency by shifting to amino acid catabolism as an energy source^41^. Amino acids are also used in protein biosynthesis and as precursors of nucleotides, lipids and cell wall components. In the present study, most of the amino acids start to be consumed when preferred carbon sources become limited. In contrast, glycine is consumed before glucose depletion, suggesting that it plays an essential role in maintaining SE growth (Supplementary Table S2). Glycine participates in pentaglycine bridges linking peptidoglycan units of the cell wall^48^. Similarly, most of the other measured gluconeogenic amino acids, which can be converted to glucose through gluconeogenesis, are depleted (alanine, aspartate, asparagine, threonine and serine) or more consumed (glutamate and methionine) by both strains at pH 7.4. For the acidic pH, this uptake seems to be delayed, probably explaining the lower biomass achieved at this pH (Fig. 3e). The same effect on this type of amino acids was reported for *S. aureus* at pH 7.4^49^, which are redirected to pyruvate synthesis to sustain biomass growth. The gradual consumption of amino acids reported here at pH 5.5, was attributed to protein synthesis and not to catabolic processes^49^. Other reports corroborate that a more active amino acid catabolism is found at blood-mimicking pH than at skin-mimicking pH^18,47^.

Although several similarities were found for the amino acid intake/secretion described here with that of *S. aureus*^29,47^, some differences were also depicted. In particular, significant uptake of lysine and leucine or secretion of tyrosine that were not noticed for the two SE strains at pH 5.5 or 7.4. In opposition, at pH 5.5 tyrosine is consumed by ICE25 (Supplementary Fig. S5). It was proposed that *S. aureus* uses lysine to balance cytoplasm pH^47^, however SE could resort to other mechanisms as suggested above.

### Nucleosides and uracil uptake are associated with a higher biomass yield at blood pH

The measured nucleosides, uridine and adenosine, are completely depleted at mid/late-exponential phase at both pH conditions and for both strains (Fig. 3f). This could suggest that at initial growth phases, SE imports nucleosides needed for cellular division or protein and peptidoglycan biosynthesis from the media instead of resorting to *de novo* synthesis. However, differential proteomics between the two pH conditions has shown that both strains have the purine and pyrimidine metabolism enhanced at a mid-exponential phase when grown at blood pH^18^. These data suggest a higher requirement for nitrogenous bases at blood pH. Interestingly, uracil is completely depleted during the exponential phase, and it is exported to the media after the late exponential phase, except by 19N at skin pH, which stops secretion at the beginning of the stationary phase (Fig. 3f). Extracellular accumulation of uracil in the stationary phase was previously reported for *S. aureus* COL^29^ and for other bacterial species^30^. It was suggested that the export of uracil by gut bacteria causes inflammation required for pathogen clearance and host survival^50^. Although poorly understood, uracil further regulates TCA cycle in multiresistant *S. aureus*^51^ and modulates virulence, biofilm formation and quorum sensing in *P. aeruginosa*^52^. Although the export of uracil in the stationary phase is a common trend in bacteria further studies are required to clarify its role. Our data suggest that both strains uptake nucleosides from the media for bacterial growth during the exponential phase.

### Regulation of Carbon Central Metabolism and Virulence factors

The catabolite control protein A (CcpA), a trans-acting primary regulator of carbon catabolite repression, increases the expression of genes encoding glycolytic enzymes and decreases the expression of genes encoding TCA cycle enzymes upon glucose addition^53^. CcpA is responsible for changing the carbon flux through these central metabolic pathways. Additionally, CcpA is involved in the expression of genes associated with biofilm production (*ica* and *cidA*) and other virulence factors through the regulation of RNAIII^54^ (Fig. 5a). Deletion of *ccpA* in SE strains acts as a positive effector of biofilm production and *icaABCD* transcription and as repressor of TCA cycle activity^16^. Our results suggest reduced activity of the TCA cycle during the exponential phase for both strains and pH conditions (see chapter “TCA cycle is more active at pH 7.4 for 19N”). This led us to hypothesise higher CcpA activity for this growth phase, which could have impact on the production of virulence factors.

**Figure 5.**
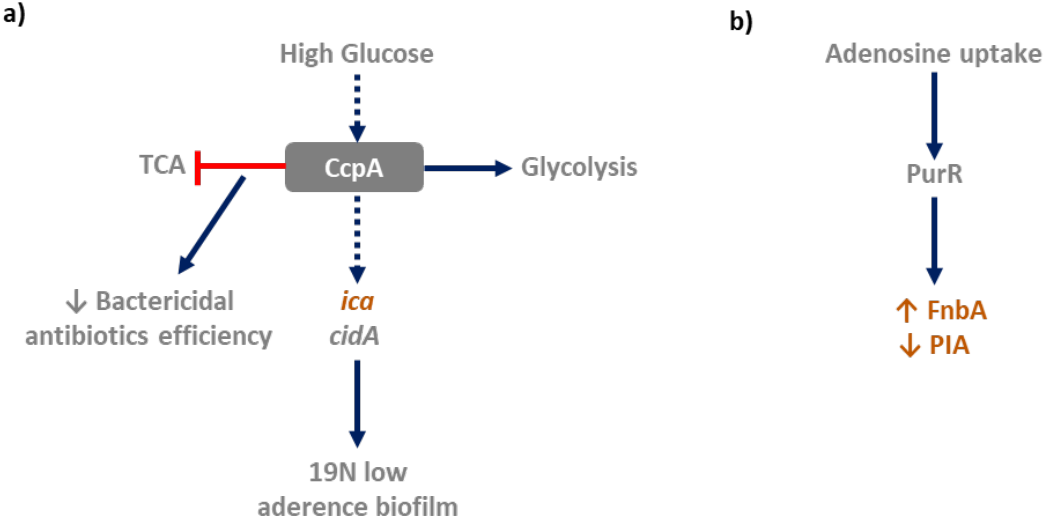
Regulation of Carbon Central Metabolism and Virulence factors associated with (A) TCA cycle regulation and (B) purine biosynthesis. Dashed arrows correspond to indirect connections, and the blocked arrow to pathway repression. Vertical arrows in the text boxes correspond to level increase (↑) or decrease (↓). Orange is used for ICE25-specific processes

It was reported for SE that PIA, a key component of biofilm that also contributes to immune evasion, is produced when the TCA cycle is repressed^16, 55^. A lower TCA cycle activity leads to decreased NADH/NAD^+^ ratio, altering the cellular redox state resulting in the derepression of *icaABDC* operon through some of its regulators^56^and, concomitantly, in the production of PIA and biofilm. PIA biosynthetic machinery encoded in the *ica* locus is only present in the ICE25 genome. Potentially, this strain can synthesize PIA at both pHs mainly during the exponential phase, while 19N may rely on other processes, which should be investigated in future studies. Nevertheless, *cidA*, present in the genome of both strains, which increases murein hydrolases and autolysis activity, could work as an important regulator of programmed cell death, which was previously associated with biofilm development through eDNA release^57^. Similarly, to what was determined for SE strain 1585WT deficient in *ica*, as for 19N, biofilms formed in human plasma are loosely attached consisting of suspended aggregates^58^. Based on these facts, it is expected that ICE25 can behave as a more virulent strain than 19N, due to the production of a more protective biofilm due to higher adhesion. Moreover, associated with decreased TCA cycle activity is the reduction of the common killing effect of most bactericidal antibiotics since low levels of superoxide and hydroxyl radicals are formed due to low oxidation of NADH via electron transport chain^59^. This suggests that during the exponential phase, for which TCA cycle flow is reduced, antibiotics efficiency is lower, which, together with the ICE25 potential to produce stronger biofilms may be advantageous for this strain proliferation in bloodstream conditions.

We found that adenosine levels decrease during the lag phase, reaching exhaustion before the mid-exponential at both pH and for both SE strains. Consumption of extracellular adenosine is mostly associated with its intake and intracellular use, suggesting that till the mid-exponential phase the purine biosynthesis is repressed. The purine biosynthesis repressor gene, *purR*, was also shown to be involved in *S. aureus* virulence modulation^60^. A *purR* defective *S. aureus* mutant leads to the overexpression of fibronectin-binding protein A, FnbA, increasing bacterial burden^60^. It was also reported that a SE *purR* mutant generated low PIA levels and had no biofilm production inducing low virulence^66^. These associations (Fig. 5b), only relevant for ICE25 since the 19N genome does not include genes homologous to *fnbA* or *ica*, suggest a contribution to higher virulence of the first strain if a higher expression of the *fnbA* gene is confirmed.

Compared to pH 5.5, at pH 7.4 we have determined for both strains, a higher glutamate import. Additionally, for their respective clonal lineages a lower production of biofilm was measured at pH 7.4^7^. *Shibamura-Fujiogi et al* have reported that the intake of exogenous glutamate by multiresistant *S. aureus* can inhibit biofilm formation^66^. Moreover, sodium/glutamate symporter (*gltS)* transcript levels were found to increase at pH 7.4 relative to pH 5.5 in *S. aureus*^24.^ All these results lead us to propose that at blood pH there is a reduction in biofilm production for which can contribute lower glutamate import restrictions. Although not directly evaluated in this study, and based on the available literature^16,26,59,60^, the exometabolites patterns determined in this study may be associated with virulence by metabolic processes such as TCA cycle regulation, purine biosynthesis and glutamate uptake. Naturally, raised hypotheses lack further evaluation. Both SE strains seem to present higher pathogenicity at the exponential phase, when TCA is repressed and purine biosynthesis is not repressed. This is especially important for ICE25 which can produce higher adherence biofilm than 19N. Preventing SE infections may not need bactericides but be based on active products that hold the bacteria in the lag phase.

## Conclusions

Upon infection, there is a cross-talk between the host and pathogen that implies adjustments in their metabolism^41^. This fact justifies comparative metabolomic studies for strains with diverse infection abilities in various environments. To the best of our knowledge, this is the first report of a comparative metabolic dynamics between SE strains belonging to A/C and B clonal lineages at two relevant biological pH conditions, those of skin and blood. At skin pH (pH 5.5), TCA and amino acid uptake are more active for 19N. More evident at this pH is the concomitant accumulation of extracellular uracil by ICE25, which can contribute to regulate the TCA cycle and, consequently, virulence. At blood pH (pH 7.4), formate is depleted from the extracellular media by the B strain, due to the presence of a formate dehydrogenase specific from this clonal lineage B, probably working as an electron donor for energy production. As shown for other bacterial species, formate accumulation in ICE25 media may constitute a virulence factor with several impacts on host invasion. An intense secretion of lactate by this strain and its lower efficiency in reverting media acidification can potentiate this impact. 19N seems better adapted to use diverse carbon sources, including ethanol and acetate. TCA cycle regulation, purine biosynthesis and glutamate uptake could be for both SE strains associated with virulence, particularly biofilm production, especially for ICE25, which is able to produce higher adherence biofilm.

Apart from the above-mentioned specificities for each strain, both strains share some metabolic patterns. SE strains preferentially utilise monosaccharides and sucrose as carbon sources. Fructose uptake was favoured over glucose at acidic pH levels since its oxidation constitutes a shortcut to glycolysis. The consumption of different saccharides is probably regulated by extracellular pH. After saccharide depletion, and independently of the media pH, fumarate, pyruvate, lactate and 2,3-BD were used as carbon sources. At pH 7.4, there is a higher demand for saccharides and amino acids together with a higher accumulation of TCA cycle intermediates and overflow metabolism products, suggesting a higher metabolic activity that may be associated with the higher biomass produced up to the stationary phase. Survival of both strains seems to be at risk below pH 5, therefore, a relevant metabolic effort is put into preventing extensive media acidification, namely through the secretion of neutral components, quaternary ammonium compounds (betaine and choline) and ammonium.

Furthermore, the dynamics of SE’s exometabolite profiles were shown to be a good proxy for complementing the understanding of the active intracellular processes and their relevance during bacterial growth at diverse pH conditions.

## Methods

Details of the methods used can be found in the Supplementary Information file.

### SE growth at pH-mimicking skin and blood environments

SE growth assays were performed for 19N and ICE25 strains at pH mimicking SE skin colonisation and blood infection (pH 5.5 and 7.4, respectively) in three independent replicates following the protocol described in ^18^. The cultures of each strain were incubated until the strains reached the stationary phase. Hourly, an 800 µL aliquot was collected to measure OD_600nm_ and the medium was separated from the cells by centrifugation at 5000 *g*, 5 min and 4°C. The supernatants were kept at −20°C for a maximum of 3 months until NMR analysis. The extracellular pH was measured in the supernatant.

### ^1^H-NMR spectroscopy of *SE* growth media

A total of eight time-points were analysed: t_0h_ collected immediately after inoculation, t_2h-6h_ hourly during the exponential phase, t_8h_ and t_10h_ during the late exponential and beginning of the stationary phase. After thawing, 540 µL of the media samples were transferred to 5 mm NMR tubes and 60 µL of the NMR buffer (1 M phosphate potassium buffer, 2 mM sodium azide, 3.22 mM 3-(Trimethylsilyl)propionic-2,2,3,3-d_4_ acid sodium salt (TSP), pH 7.0) was added. NMR spectra were acquired on a Bruker Avance II+ 800 MHz spectrometer equipped with a 5 mm TXI-Z H/C/N/-D probe at 298.15 K, as previously described^18^. Chenomx Suite 8 was used for metabolites assignment and absolute quantification. The areas of the unassigned resonances (Unks) were integrated with NMRProcflow^63^. Low intensity Unks peaks led to no corresponding peaks in TOCSY. To check whether those Unks belong to the same metabolite, Pearson correlation between them was performed. If a strong correlation between two Unks was observed (coefficient > 0.9), they were considered as possibly belonging to the same metabolite (although allowing for concomitant biochemical relationships). Areas of the unassigned signals were normalised to the TSP area and analysed as described below. NMR raw data was deposited into the NIH Common Fund’s National Metabolomics Data Repository (NMDR) via the Metabolomics Workbench Project ID PR001917 and is available at http://dx.doi.org/10.21228/M8XH8G.

### Analysis of metabolite profiles

Before NMR profiling, the time points considered for analysis were adjusted for the growth phases by translating the curves to obtain superimposable exponential phases. For pH 5.5, the growth time of replicate one was shifted by minus 2 and 3 hours for 19N and ICE25 strains, respectively. For pH 7.4, replicates one and three were shifted by minus 2 and 1 hour, only for strain 19N. Biological replicates of ICE25 grown at pH 7.4 were shown to be highly superimposable.

A permutation filter was used to assess whether metabolite levels had changed significantly over time within each experimental condition^21^. Briefly, for each metabolic feature (identified metabolite or Unks) an estimate of the total variation (*SDsum*_*true*_) was calculated as in equation 1:

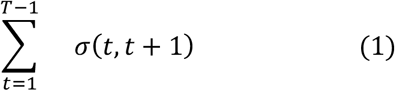

where *T* is the number of time points and *σ*(*t, t* + 1) is the standard deviation of the variable concentration at two consecutive time points, t and t+1. Then 1000 permutations were performed in the time axis, and the corresponding variances, *SDsum*_*perm*_, were estimated using (1). A metabolic feature profile was considered significantly changed if: *SDsum*_*true*_< 0.25 *quantile of SDsum*_*perm*_.

The correlation of metabolic features with medium pH was determined based on the Pearson coefficient for each experimental condition, considering the time intervals before and after the minimum medium pH value.

Metabolic feature concentrations were mean centred and divided by standard deviation (Z-scores). A K-means clustering with Dynamic Time Warping (DTW) was performed to identify similar profiles in each pH condition^64^. The optimal number of clusters was chosen based on the Elbow Method^65^. K-means clustering was performed 24 times to access the most frequent cluster of each one of the metabolic features. Principal component analysis (PCA) of the time course of each strain’s metabolic features was performed to evaluate if the dynamics were similar between strains within each pH.

All data analyses described were performed in a Python3 environment using in-house scripts and are available through GitHub access at https://github.com/eccmorais/Exometabolomics_Sepidermidis.git.

To search for possible transporters related to the translocation of the measured exometabolites and metabolic pathways in which those may be involved, the BioCyc database for the reference strain, SE RP62A (version 27.5) was accessed^66^.

## Author contributions

E.M., L.G.G., M.M. and A.V.C. contributed to the study conception and design. E.M. performed material preparation, data collection and analysis All authors were involved in data interpretation. E.M. wrote the first draft of the manuscript, and all authors commented on and contributed to the manuscript’s writing. All authors reviewed the manuscript.

## Data availability statement

NMR raw data was deposited into the NIH Common Fund’s National Metabolomics Data Repository (NMDR) via the Metabolomics Workbench Project ID PR001917 and is available at http://dx.doi.org/10.21228/M8XH8G.

