## Supplementary material for "Dynamic exometabolomics reveals metabolic adaptations of *Staphylococcus epidermidis* to pH-mimicking skin and bloodstream"

Supplementary Information

### Detailed methods

#### SE growth at pH-mimicking skin and blood environments

SE growth assays were performed for 19N and ICE25 strains at pH mimicking SE skin colonisation and blood infection (pH 5.5 and 7.4, respectively) in three independent replicates. Briefly, strains were inoculated from the glycerol stock in Tryptic Soy Agar (TSA, BD Biosciences, pH 7.4) overnight, and a single colony was picked and incubated overnight again in a new TSA plate (pH 7.4). A colony of each strain was picked and incubated in Tryptic Soy Broth (TSB, BD Biosciences, pH 7.4 or pH 5.5) at 37°C, 225 rpm overnight. The $OD_{600nm}$ was adjusted to 0.06 (final volume 100 mL). The cultures were incubated at 37°C, 225 rpm until the strains reached the stationary phase. The volume of the culture medium was kept at 36% (90 ml/250 ml) of the flask total volume to ensure a good aeration. Hourly, an 800 µL aliquot was collected to measure $OD_{600nm}$ and the medium was separated from the cells by centrifugation at 5000 *g*, 5 min and 4°C. The supernatants were kept at -20°C until NMR analysis for a maximum of 3 months. The extracellular pH was measured in the supernatant.

#### $^1$H-NMR spectroscopy of *SE* growth media

A total of eight time-points were analysed: $t_{0h}$ collected immediately after inoculation, $t_{2h-6h}$ hourly during the exponential phase, $t_{8h}$ and $t_{10h}$ during the late exponential and beginning of the stationary phase. After thawing, 540 µL of the media samples were transferred to 5 mm NMR tubes and 60 µL of the NMR buffer (1 M phosphate potassium buffer, 2 mM sodium azide, 3.22 mM 3-(Trimethylsilyl)propionic-2,2,3,3-$d_4$ acid sodium salt (TSP), pH 7.0) was added. NMR spectra were acquired on a Bruker Avance II+ 800 MHz spectrometer equipped with a 5 mm TXI-Z H/C/N/-D probe at 298.15 K. 1D $^1$H-NMR spectra were obtained using the *noesygppr1d* pulse program with the following parameter: 64 scans; 4 s of relaxation delay; 10 ms of mixing time; 16025.641 Hz spectral width; 128 k points of the size of Free Induction Decay signal (FID). Each FID was multiplied by a 1.0 Hz exponential line broadening function, followed by Fourier Transformation. All spectra were manually phased, baseline corrected and adjusted to the TSP chemical shift at 0.00 ppm. Chenomx Suite 8 was used for metabolites assignment and absolute quantification. The areas of the unassigned resonances (Unks) were integrated with NMRProcflow (Jacob et al., 2017). Low intensity Unks peaks led to no corresponding peaks in TOCSY. To check whether those Unks belong to the same metabolite, Pearson correlation between them was performed. If a strong correlation between two Unks was observed (coefficient > 0.9), they were considered as possibly belonging to the same metabolite (although allowing for concomitant biochemical relationships). Areas of the unassigned signals were normalised to the TSP area and analysed as described below. Metabolomic data is available at the NIH Common Fund's National Metabolomics Data Repository (NMDR) via the Metabolomics Workbench Project ID PR001917 (doi: http://dx.doi.org/10.21228/M8XH8G).

#### Evaluation of ICE25 intracellular metabolome after medium sterilization by autoclave or filtration

The growth of ICE25 was carried out O/N at 37 °C on Tryptic Soy Agar plates (TSA, BactoTM). A single colony was used to pre-inoculate the TSB medium adjusted to pH 5.5 and 7.4. Two sterilization conditions were tested: autoclaving at 120°C (A) or filtration on a laminar flow chamber (F) for each pH.

The pre-inoculum was adjusted to a cellular density of $1.5x10^8$ CFU/mL ($OD_{600}$=0.06) to inoculate 90 mL of TSB in 250 mL erlenmeyers. ICE25 was grown O/N at 37 °C and 225 rpm. The bacterial cells were harvested at the mid-exponential phase by centrifugation at 5000 *g* for 5 min. Five biological replicates of independent growth assays were prepared for each experimental condition.

Cell pellets were resuspended in 20 mM phosphate buffer pH 7.2-7.4 to adjust the final $OD_{600}$ to 20. Metabolites were extracted with methanol/chloroform/water (5:5:1) and dried in a SpeedVac. To prepare the samples for NMR, 750 µL of phosphate buffer (33 mM, pH 7.0 in $D_2O$ with 2 mM of sodium azide) with 0.21 mM of 3-(trimethylsilyl)propionic-2,2,3,3-$d_4$ (TSP) was added, and 600 µL was transferred to 5 mm NMR tubes.

NMR spectra were acquired on a Bruker Avance II+ 800 MHz spectrometer equipped with a 5 mm TXI-Z H/C/N/-D probe at 298.15 K. 1D $^{1}H$-NMR spectra were obtained using a *noesygppr1d* pulse program.

Spectra were processed with Topspin 4.0 and with NMRProcFlow (https://nmrprocflow.org/) that it was used to create the spectra buckets used for the statistical analysis on Metaboanalyst (https://www.metaboanalyst.ca/). Multivariate statistical analysis, Principal Component Analysis (PCA) and Partial Latent Square- Discriminate Analysis (PLS-DA), were performed.

**Supplementary Figure S1. Twenty metabolic features levels are significantly different at $t_{0h}$ between pH conditions.** Box-plots are organized from features increased at pH 7.4 (negative R) to features increased at pH 5.5 (positive R). R, corresponds to the ratio of the metabolic feature areas between pH 5.5 and pH 7.4 in percentage and considering 0% as no changes between pH (average area at pH 5.5 /average area at pH 7.4)*100-100)

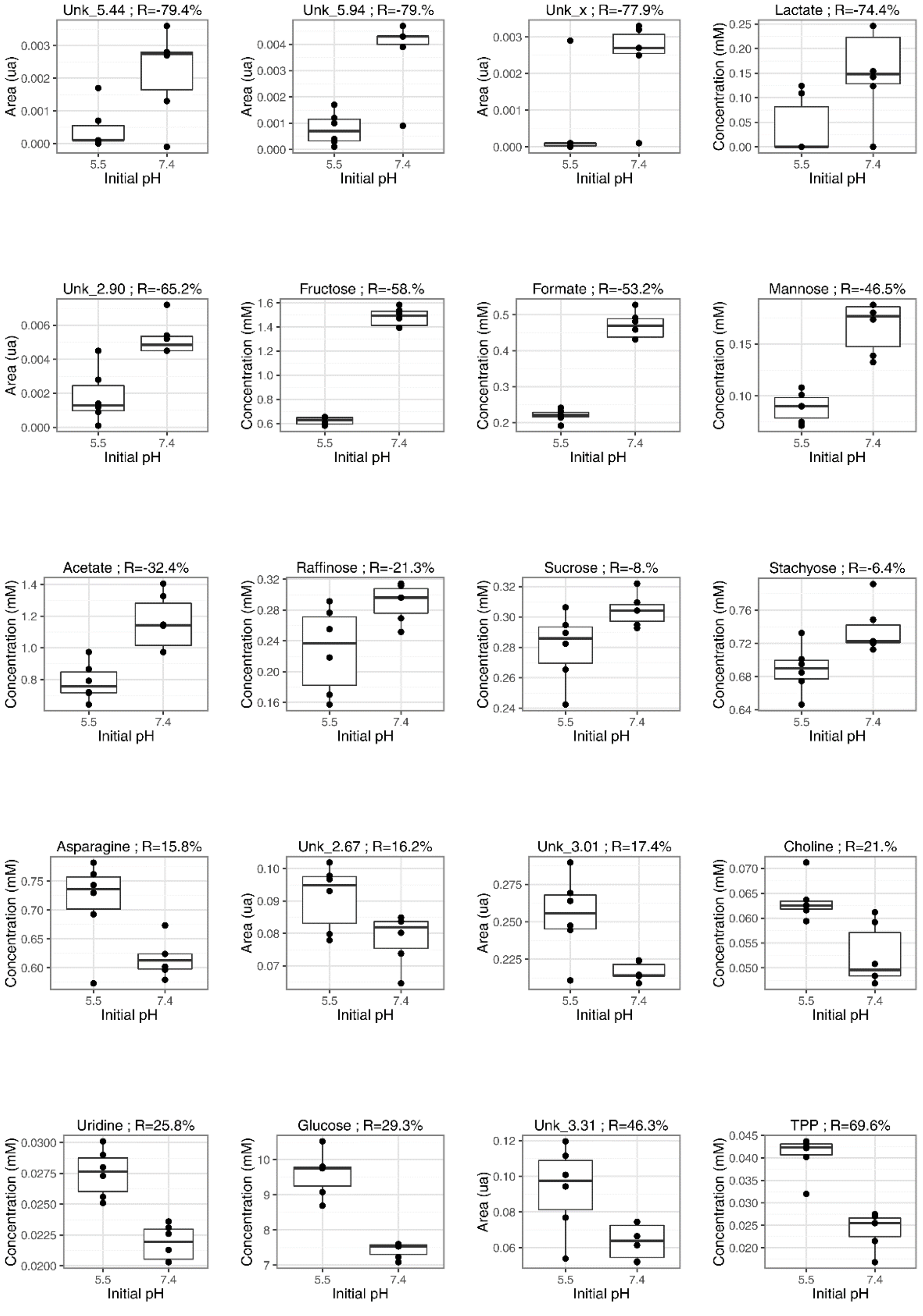

**Supplementary Figure S2. TSB media glucose levels after the sterilization process are significantly diminished at pH 7.4.** $^1$H-NMR resonance of anomeric glucose duplet at 5.25 ppm for TSB media adjusted to pH 7.4 and 5.5, before and after the sterilization process

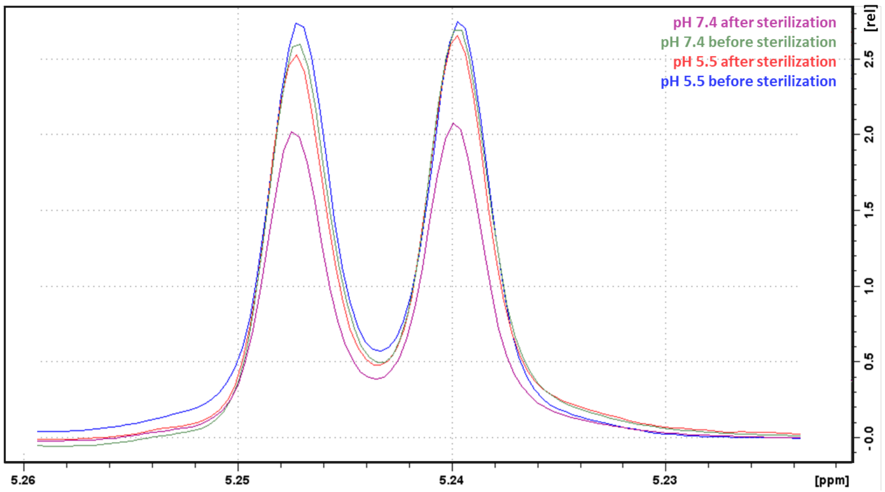

**Supplementary Figure S3. Effect of sterilization method on the ICE25 endometabolome.** A) PCA scores plot of all the conditions: autoclaved media at pH 5.5 (5A) and at PH 7.4 (7A); filtrated media at pH 5.5 (5F) and at pH 7.4 (7F). B) PLS-DA score plot, Q2 and permutations test for comparison of autoclaved and filtrated media at pH 7.4, showing that the model it is not valid. C) PLS-DA score plot, Q2 and permutations test for comparison of autoclaved and filtrated media at pH 5.5, showing that the model it is not valid.

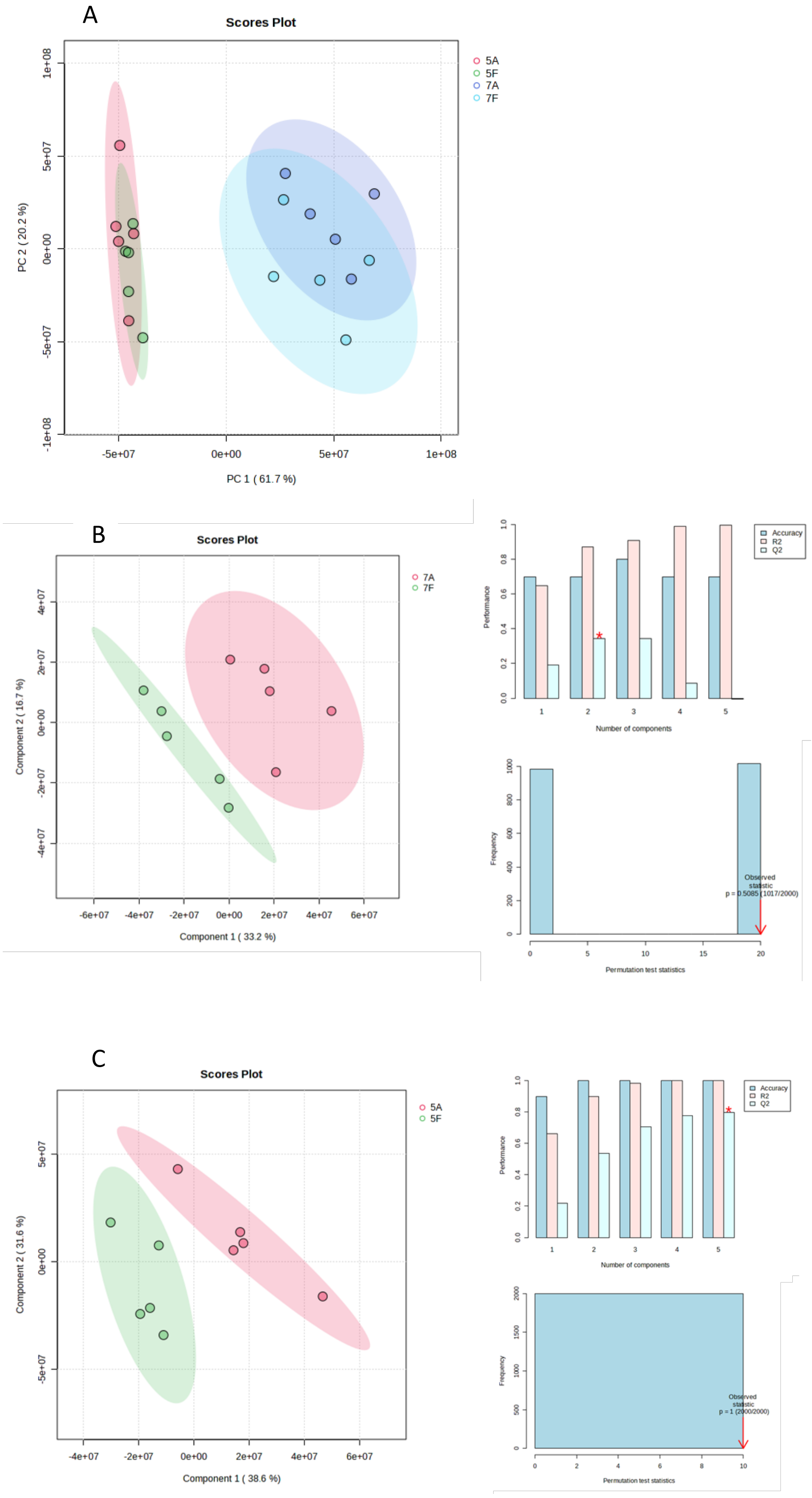

**Supplementary Figure S4. Extracellular OD and pH measurements without time adjustment.** Biomass (OD, optical density at 600 nm, left y-axis) and extracellular pH (right y-axis) average curves for 19N and ICE25 strains grown in TSB at initial pH of 5.5 and 7.4. Filled and dotted lines correspond to OD and extracellular pH measurements with standard deviation in light grey (n=3)

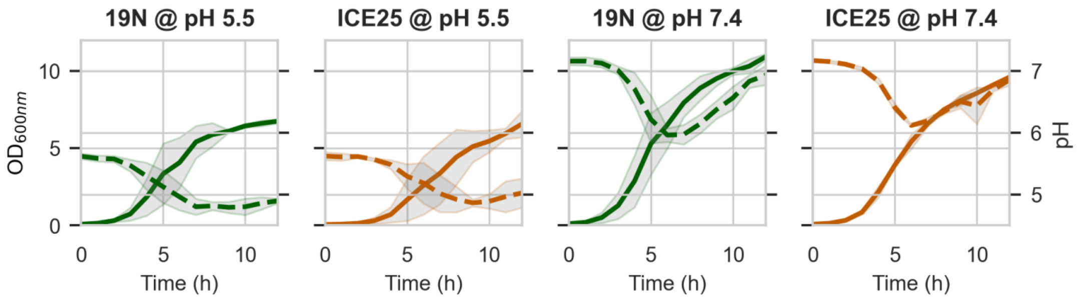

**Supplementary Figure S5. Temporal dynamics of the metabolic features.** Solid lines correspond to the variation of metabolic feature levels for 19N and ICE25 strains by biological replicate, in green and orange (left y-axis), respectively. Dashed lines represent the bacterial growth curve

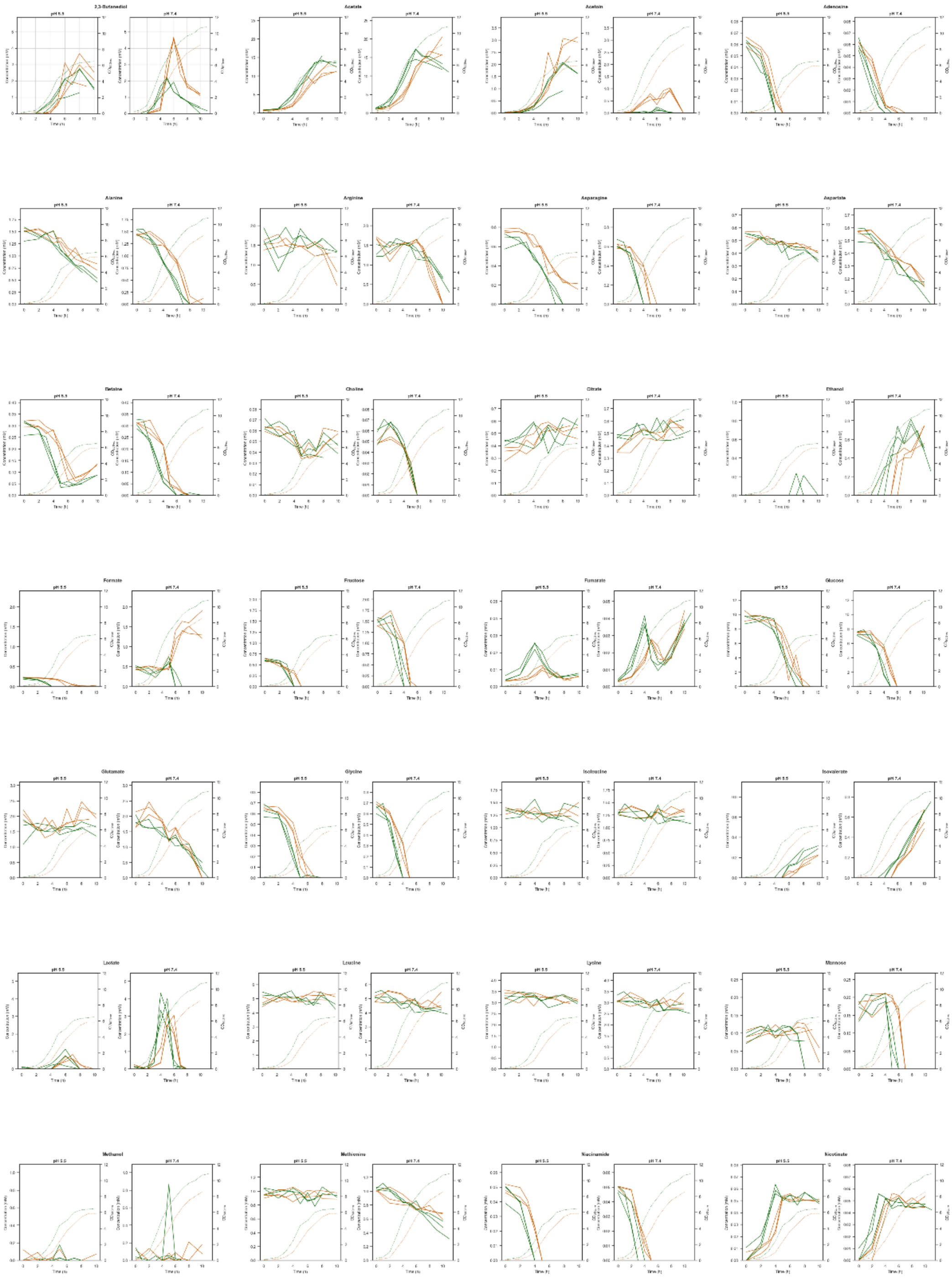

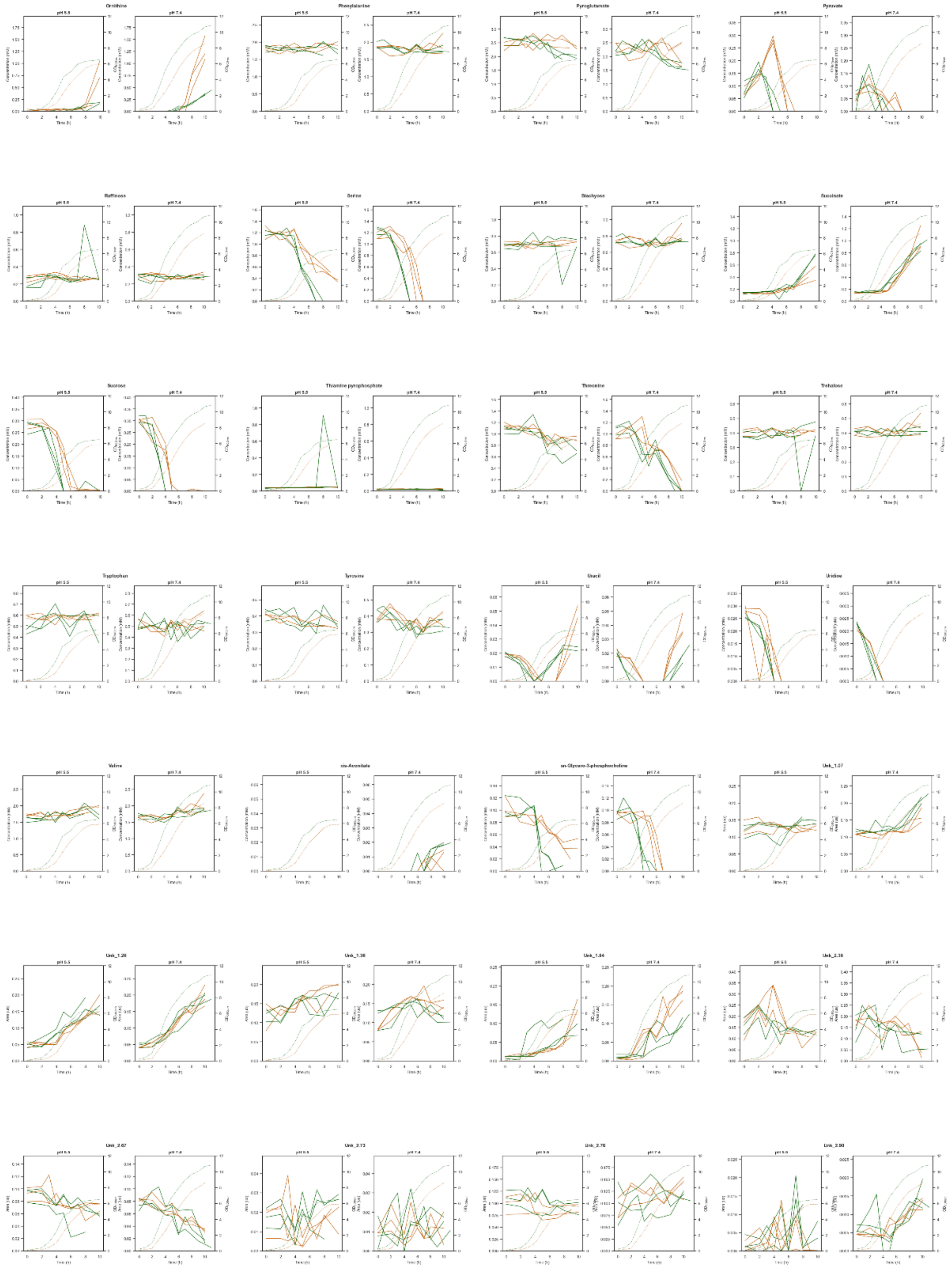
Ornithine
Phenylalanine
Pyroglutamate
Pyruvate
Raffinose
Serine
Stachyose
Succinate
Sucrose
Thiamine pyrophosphate
Threonine
Trehalose
Tryptophan
Tyrosine
Uracil
Uridine
Valine
cis-Aconitate
sn-Glycero-3-phosphocholine

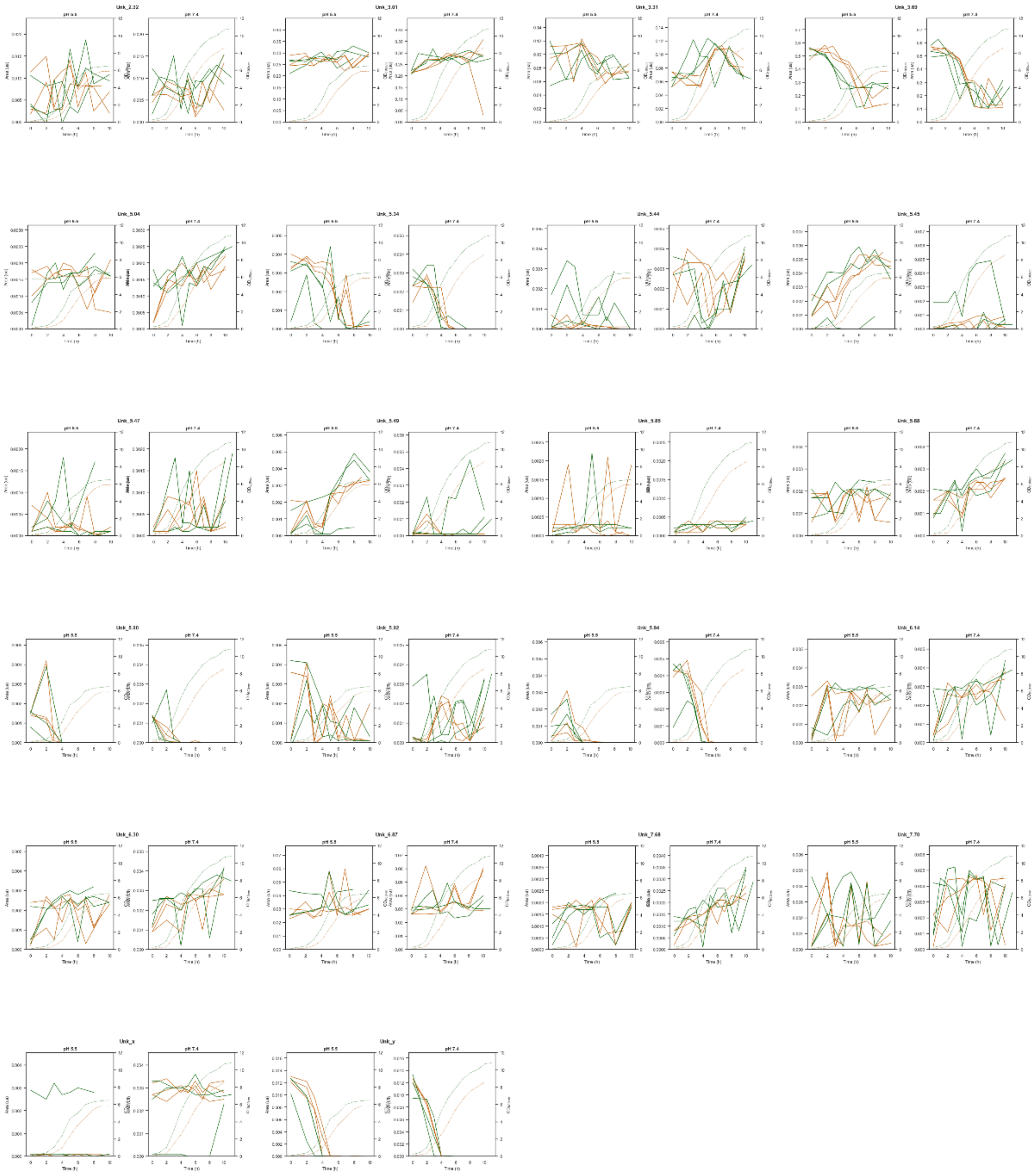

**Supplementary Figure S6. Correlation of metabolite profiles with pH dynamics on the experimental condition.** Correlation of metabolic features with extracellular pH for each experimental condition and each pH variation. Pearson correlation coefficient and associated *p*-values were determined. The gradient from blue to red represents the values ranging from a strong negative to a positive correlation (values ranging from -1 to 1 ). Some metabolic feature names are abbreviated (see Table S2)

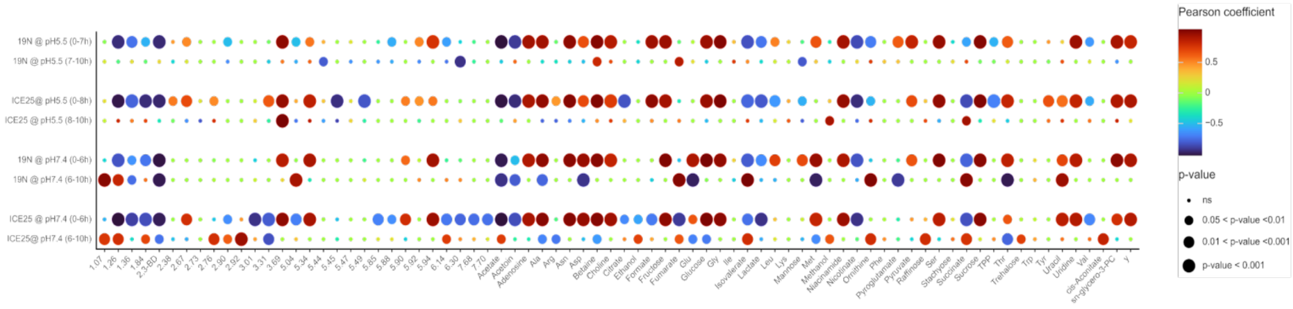
